# Inference of genome 3D architecture by modeling overdispersion of Hi-C data

**DOI:** 10.1101/2021.02.04.429864

**Authors:** Nelle Varoquaux, William S. Noble, Jean-Philippe Vert

## Abstract

We address the challenge of inferring a consensus 3D model of genome architecture from Hi-C data. Existing approaches most often rely on a two step algorithm: first convert the contact counts into distances, then optimize an objective function akin to multidimensional scaling (MDS) to infer a 3D model. Other approaches use a maximum likelihood approach, modeling the contact counts between two loci as a Poisson random variable whose intensity is a decreasing function of the distance between them. However, a Poisson model of contact counts implies that the variance of the data is equal to the mean, a relationship that is often too restrictive to properly model count data.

We first confirm the presence of overdispersion in several real Hi-C data sets, and we show that the overdispersion arises even in simulated data sets. We then propose a new model, called Pastis-NB, where we replace the Poisson model of contact counts by a negative binomial one, which is parametrized by a mean and a separate dispersion parameter. The dispersion parameter allows the variance to be adjusted independently from the mean, thus better modeling overdispersed data. We compare the results of Pastis-NB to those of several previously published algorithms: three MDS-based methods (ShRec3D, ChromSDE, and Pastis-MDS) and a statistical methods based on a Poisson model of the data (Pastis-PM). We show that the negative binomial inference yields more accurate structures on simulated data, and more robust structures than other models across real Hi-C replicates and across different resolutions.

A Python implementation of Pastis-NB is available at https://github.com/hiclib/pastis under the BSD license

Supplementary information is available at https://nellev.github.io/pastisnb/

## 1. INTRODUCTION

DNA *in vivo* is folded in three dimensions, and this 3D structure plays an important role in many biological functions, including gene regulation, DNA replication, and DNA repair^8,9,30,33^. Chromosome conformation capture methods, coupled with next-generation sequencing, allow researchers to probe the three-dimensional structure of chromosomes within the nucleus^21^. These techniques, which we broadly refer to as “Hi-C,” rely on crosslinking, digesting, ligating, and paired-end sequencing of DNA to identify physical interactions between pairs of loci. Hi-C techniques provide a genome-wide *contact map*, a matrix indicating the contact frequency between pairs of loci. This matrix can be used to analyze the three-dimensional structure of the genome. However, despite extensive research, inferring a three-dimensional model from this contact map remains a fundamental problem.

Methods to infer the 3D structure of the genome broadly fall into two categories: *ensemble* approaches which infer a population of structures^15,18,29^, and *consensus* approaches which yield a single model that summarizes the contact count data^2,4,34,36,39^. The former approach is more biologically accurate because the population of models better reflects the diversity of structures present in a population of cells. However, interpretation of the resulting models is challenging, and one often has to fall back to a single structure or a few structures that best represent the population of structures. Validation of ensemble models, even on simulated data, can also be challenging. On the other hand, a consensus model summarizes the hallmarks of genome architecture, can easily be visually inspected and analysed, and can be integrated in a straightforward manner with other data sources, such as gene expression, methylation, and histone modifications, which are also ensemble based. In this work, therefore, we focus on inferring a consensus model of the 3D genome architecture.

Consensus approaches model chromosomes as chains of beads, minimizing a cost function that aims to produce a model as consistent with the data as possible. In addition, the optimization is sometimes constrained to include prior knowledge about the 3D structure: the size and shape of the nucleus, distance constraints between between pairs of adjacent loci, etc. Some methods first convert contact counts into *wish distances*, either through a biophysically motivated counts-to-distances mapping or through *ad hoc* conversions. They then use multidimensional scaling (MDS) methods such as metric MDS^10,34^, weighted metric MDS^2^, non-metric MDS^4^, or classical MDS (most commonly known as PCA)^19^. These methods rely on arbitrary loss functions.

In previous work, we introduced a consensus method called “Pastis,” based on a statistical model of contact count data, where the 3D structure is the latent variable and the inference of the consensus 3D model is formulated as a maximum likelihood problem^36^. A natural statistical model for count data is the Poisson distribution, which has a single parameter, the mean *µ*, from which all its other properties (including the variance) are derived. Pastis relies on such a Poisson model of the interaction frequencies, the intensity of which decreases with the increasing spatial distance between the pair of loci. We further extended this Poisson modeling to allow for inference of diploid structures^7^. We refer to this method below as “Pastis-PM” (“PM” standing for “Poisson model”).

However, a Poisson model of contact counts implies that the variance of the data is equal to the mean, an assumption that is sometimes too restrictive to properly model the data. For instance, Nagalakshmi *et al*. ^23^ and Robinson and Smyth ^26^ show that for RNA-seq, this assumption is not justified, because the variance in the data is larger than the mean, leading to to an overdispersion problem. To alleviate the overdispersion, Robinson *et al*. ^28^ suggest modeling RNA-seq data with a negative binomial distribution, which is parameterized with two parameters: the mean *µ* and the variance *σ*. Modeling Hi-C contact count data with a negative binomial model is not new. Jin *et al*. ^17^ and Carty *et al*. ^6^ used this approach to assign statistical confidence estimates to observed contacts, Hu *et al*. ^14^ to normalize the data, and Lévy-Leduc *et al*. ^20^ to find topologically associated domains.

In this work, we explore methods that apply a similar generalization—from Poisson to negative binomial— in the context of a model for inferring genome 3D structure from a Hi-C contact map. We first confirm the overdispersion on a wide variety of Hi-C datasets, from very small (*S. cerevisiae*) to large genomes (human). We then compare our method based on a negative binomial model for Hi-C count data, which we call Pastis-NB, to MDS-based methods (chromSDE, ShRec3D, and Pastis-MDS) and to the Poisson model-based Pastis-PM. We first demonstrate that Pastis-NB recovers the most accurate results, in particular in low coverage settings. We then study how well the different methods perform when provided with an incorrect mapping between contact counts and Euclidean distances, a setting where Pastis-NB also outperforms other methods. Finally, we show that Pastis-NB model yields more stable structures across Hi-C replicates and across resolutions.

### II. APPROACH

We model each chromosome as a series of *n* beads, where each bead corresponds to a specific genomic window, which we refer to as a *locus*. We aim to infer the coordinate matrix **X** = (*x*_1_, *x*_2_,…, *x*_*n*_) ∈ ℝ^3*×n*^, where *x*_*i*_ corresponds to the 3D coordinates of bead *i*. We denote by *d*_*ij*_ = *x*_*i*_ *x*_*j*_ the Euclidean distance between beads *i* and *j*. Hi-C contact counts can be summarized as an *n*-by-*n* matrix **c** in which rows and columns correspond to loci, and each entry *c*_*ij*_ is an integer, called a *contact count*, corresponding to the number of times loci *i* and *j* are observed in contact. This matrix is by construction square and symmetric. Since raw contact counts are known to be biased by loci-specific multiplicative factors, we apply ICE normalization^16^ to estimate a bias vector *b* = (*b*_1_,…, *b*_*n*_) ∈ ℝ^*n*^, where *b*_*i*_ is the bias factor for locus *i*. The normalized contact count matrix **c**^*N*^ is then defined as the matrix of normalized counts 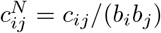. See Supplementary Materials **??** for more details.

### II.A. Statistical model

We model the raw contact counts *c*_*ij*_ as realizations of independent negative binomial random variables

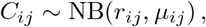

where *r*_*ij*_ is the dispersion parameter and *µ*_*ij*_ is the mean of the negative binomial distribution between loci *i* and *j*. Like in the Pastis model we parameterize the mean count value *µ*_*ij*_ as a decreasing function of the distance between beads *i* and 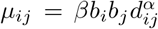, with parameters *β >* 0 and *α <* 0. *β* can be thought of as a scaling factor—the higher the coverage of the dataset, the higher it is—while *α* characterizes how the frequency of contacts decreases with the distance. Note that this relationship is ill-posed, in the sense that the relationship between the mean of the distribution *µ*, the scaling factor *β*, and the Euclidean distances *d* is not unique. In particular, one can choose to set *β* to 1 in order to infer the 3D coordinates, and then rescale the inferred structure to reflect prior knowledge about the size of the nucleus. We thus drop the scaling parameter *β* in the rest of the derivations. In addition, we parameterize the dispersion as *r*_*ij*_ = *b*_*i*_*b*_*j*_*r*, where *r* ≥ 0 accounts for overdispersion.

The probability mass function can thus be written as:

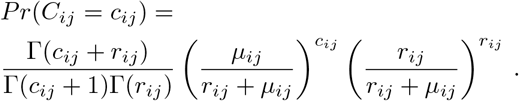

It is well-known that the variance of *C*_*ij*_ satisfies

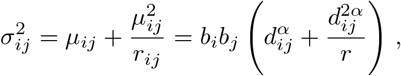

and that when *r*_*ij*_→ ∞ the negative binomial distribution tends to a Poisson distribution with intensity parameter *µ*_*ij*_. The negative binomial is thus a generalization of the Poisson distribution where the variance of the data can exceed the mean, as controlled by the dispersion parameter^24^.

### II.B. Estimating the dispersion parameters *r*_*ij*_

To estimate the dispersion parameters *r*_*ij*_, we leverage the property that the variance of an NB(*ρ, µ*) random variable is *σ*^2^ = *µ* + *µ*^2^*/ρ*; therefore, if we know *µ* and *σ*^2^, we deduce the dispersion as *ρ* = *µ*^2^*/ σ*^2^ *µ*. This implies that if a relationship *σ*^2^(*µ*) between the mean and variance of the NB distribution is known or assumed, then the dispersion parameter *ρ* is also known as a function of *µ* as *ρ*(*µ*) = *µ*^2^*/ σ*^2^(*µ*) − *µ*. We thus focus on estimating the variance as a function of the mean of the contact counts in order to infer the dispersion parameter *r*.

In the case of RNA-seq, the function *σ*^2^(*µ*) is usually estimated by fitting a weighted least squares^3^, fitting a loess^1^, or by maximizing a likelihood on empirical means and variances estimated for each gene using replicates^27^. In the case of Hi-C, relying on biological replicates is not a viable option because most studies perform at most two biological replicates, rendering the estimation of the mean and variance impossible. Instead, we estimate *σ*^2^(*µ*) by introducing additional modeling assumptions that capture properties intrinsic to Hi-C data and genome architecture. We propose a two-step method to estimate this function. First, we compute for each genomic distance *l* the empirical mean and variance of the normalized contact counts:

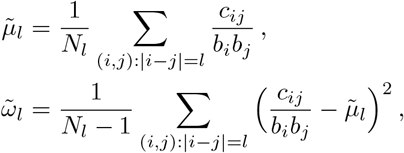

where *N*_*l*_ is the number of (*i, j*) pairs with *i j* = *l*. As shown by Yu *et al*. ^37^, 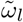 is a biased estimator of the variance and can be corrected to be unbiased as follows:

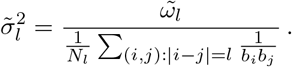

Note that in our model, estimating the mean and variance from empirical normalized counts at a given genomic distance amounts to assuming that the mean normalized count, i.e.,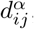, is constant for beads (*i, j*) at a given genomic distance from each other. This is obviously an assumption that we only use to get an estimate of the overdispersion parameter, but that we relax later when we optimize the 3D coordinates of each bead without constraint on their pairwise distances.

Because the empirical mean and variance may not be reliable for very long genomic distances, where the number of contact counts is small, we only compute 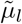 and 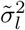 for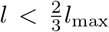, where *l*_max_ is the maximum distance between loci. We also discard genomic distances with an empirical mean or variance equal to 0.

We then proceed to estimate the dispersion parameter *r* in two steps. First, from the set of 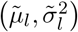 pairs, we then estimate the dispersion 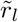 for all genomic distance as follows:

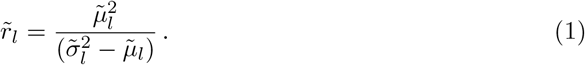

Second, from the estimates 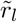 for each genomic distance, we proceed to estimating the final dispersion parameter *r* by taking the weighted average:

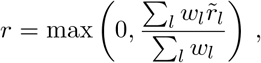

where *w*_*l*_ the number of data points used in the estimation of 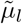 and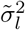. Thus, more weight is given to short genomic distances than long genomic distances.

### II.C. Estimating the 3D structure

In summary, our proposed model has three main components: (1) we model contact counts using negative binomial distributions parameterized by the mean and the dispersion parameter *C*_*ij*_∼ *NB*(*r*_*ij*_, *µ*_*ij*_); (2) we parameterize the mean as a function of the structure 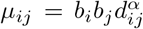; and (3) we model the dispersion parameter as *r*_*ij*_ = *b*_*i*_*b*_*j*_*r* and provide a method to estimate *r*. By combining these three components, we can write the probability of each observation:

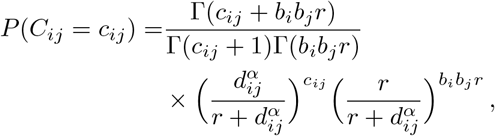

which depends on the 3D structure **X** through the pairwise distances *d*_*ij*_, and which also depends on the count-to-distance mapping parameter *α*. We then propose to jointly infer both the 3D structure **X** and the *α* parameter by maximizing likelihood:

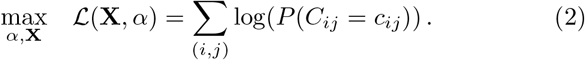

Given an observed Hi-C contact map, we solve the optimization problem of Equation 2 using the L-BFGS algorithm^5,22^ from the scipy toolbox. The optimization being non-convex, we solve it five times with random initialization to find local optima, and return the solution with the highest log-likelihood. Note that the problem in **X** is ill-posed, in the sense that the solution is defined relative to a rotation factor and a translation factor. Note also that Equation 2 is a sum over all pairs of loci (*i, j*), and may include zero counts *c*_*ij*_ = 0 which contribute to the likelihood. We illustrate in Section IV.C.1 below the benefits of keeping zero counts in the objective functions, and not filtering them.

## III. METHODS

### III.A. Simulated datasets

Because little experimental data is available to characterize the true population of 3D DNA structure, we first compare the different structural inference methods by using simulated data. We construct three ensembles of datasets with varying coverage, dispersion, and counts-to-distance mapping. All simulations use a consensus architecture obtained by running Pastis-MDS, applied to the first chromosome of the 75^th^ replicate of the KBM7 nearly haploid human cell line data from Rao *et al*. ^25^ at 100 kb.

We generate simulated datasets using the model 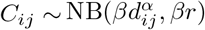. Note that we do not simulated biased data requiring ICE normalization, to focus on the architecture inference part. Since *d*_*ij*_ is given by the consensus architecture used to simulated the counts, for all pairs of beads (*i, j*), the total number of counts in the count matrix (a.k.a. coverage) is on average 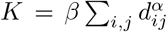; hence we control the coverage in a simulation with the *β* parameter by setting 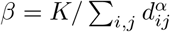.

We generate the first ensemble of 100 datasets to study the influence of coverage. We use *α* = −3, which yields a count-to-distance mapping consistent with the one obtained from polymer physics theory^12,13,21^. We vary the parameter *β* such that the expected number of reads ranges between 10% and 100% of the original dataset (10%, 20%, 30%, 40%, 50%, 60%, 70%, 80%, and 100%), and we set the dispersion parameter *r* to be the one fitted as described in Section II.B to the KBM7 Hi-C dataset (*r* = 49.9), obtaining 100 datasets by repeating each configuration 10 times with 10 random seeds.

The second collection of 100 simulated datasets is to study the influence of overdispersion. We keep *α* = −3 and set the parameter *β* such that the expected number of reads is 100% of the original dataset. We set the dispersion parameter *r* to be the one estimated on the original KBM7 contact maps multiplied by *γ*, where *γ* 0.1, 0.2,…, 1. Varying *γ* thus varies the dispersion, and the smaller the dispersion parameter is, the more overdispersed the datasets are, and thus the harder the inference is likely to be. For each set of parameters, we generate 10 datasets using 10 different random seeds, thus yielding 100 datasets.

Finally, we generate an ensemble of 70 datasets to measure how well methods perform when provided with an incorrect counts-to-distances mapping. To do so, we vary *α* ∈ {−1.5, −.2, −4., −4.5}, fix *β* for each simulation so that the number of reads is as the original dataset, keep the dispersion parameter *r* as fitted on the KBM7 Hi-C dataset, and repeat each simulation 10 times with 10 random seeds. This third ensemble of datasets enables us to compare *metric* methods, for which the counts-to-distance mapping is fixed or provided by the user *a priori*, versus *non-metric* methods, for which the counts-to-distance mapping is inferred jointly with the structure from the data.

### III.B. Real datasets

#### III.B.0.a. High-coverage in situ Hi-C from a Chronic myelogenous leukemia cell-line

We also apply our method to publicly available Hi-C data from the chronic myelogenous leukemia cell-line KBM7^25^. This cell line has the nice property of being nearly haploid: apart from chr 8 and a small part of chr 15, all chromosomes are haploid. We downloaded the first two replicates (experiment 75 and 76) and processed the data with HiC-Pro^31^ to obtain intra-chromosomal maps at 1 Mb, 500 kb, 250 kb, 100 kb, and 50 kb. We then filtered out rows and columns of the data that interact the least (See Supplementary Table S1).

#### III.B.0.b.S. cerevisiae, D. melanogaster, A. thaliana Hi-C contact counts

We downloaded publicly available whole-genome Hi-C datasets from *S. cerevisiae*^10^, *D*.*melanogaster* ^32^, and *A. thaliana*^11^. We normalize these datasets by eliminating the 4% lowest interacting loci and then applying ICE.

## IV. RESULTS

We perform a series of experiments to assess the accuracy of the different methods on simulated data and the robustness of the methods on real data. Specifically, we perform experiments on simulated data to assess the accuracy of the inferred 3D models when varying (1) the coverage of the datasets; (2) the overdispersion of the counts; (3) the counts-to-distance mapping parameter. We then assess the stability of the methods on real data on two different biological replicates; (2) across different resolution; (3) when subsampling the data; (4) on a multi-chromosome dataset, by varying the number of chromosomes included in inference. We present a summary of all results in Table I.

**Table I.** Summary of the results. We summarize here all the experiments performed, including the sections in the text where they can be found and a synopsis of the main results. The table displays the top three methods for each experiment (✓✓✓ for the best method, ✓✓ for the second best, ✓ for the third best). Pastis-NB performs better than all methods in most experiments.

|  | Simulated data |  |  | Real data |  |  |  |  |
| --- | --- | --- | --- | --- | --- | --- | --- | --- |
| | Coverage<br>(small $\beta$) | Dispersion<br>(small $\gamma$) | Counts-to-maps<br>(small $\alpha$) | Replicates<br>(LR) | Replicates<br>(HR) | Resolution | Downsampling | Multi-<br>chrom. |
| Section | 4.2.1 | 4.2.1 | 4.2.2 | 4.3.2 | 4.3.2 | 4.3.3 | 4.3.4 | 4.3.5 |
| Pastis-MDS | . | ✓✓ | . | . | . | . | . | . |
| ShRec3D | ✓✓ | . | ✓✓ | ✓ | ✓✓ | ✓✓ | ✓✓ | ✓ |
| ChromSDE | . | . | ✓ | ✓✓✓ | ✓ | ✓ | ✓ | NA |
| Pastis-PM | ✓ | ✓ | . | . | . | . | . | ✓✓ |
| Pastis-NB | ✓✓✓ | ✓✓✓ | ✓✓✓ | ✓✓ | ✓✓✓ | ✓✓✓ | ✓✓✓ | ✓✓✓ |

### IV.A. Real and simulated Hi-C data are highly overdispersed

Before diving into the comparison of the different models and algorithms, we first investigate the extent to which existing Hi-C data sets show evidence of overdispersion. Because it is rare to have several samples of the same cell line, we use the method described in Section II.B to check for the presence or absence of overdispersion in the normalized Hi-C data. We plot in Figure 1,= for different species (chr10 of the KBM7 human cell line at 100 kb, *D. melanogaster* at 10 kb, *A. thaliana* at 40 kb and *S. cerevisiae* at 10 kb), the mean vs variance relationship of normalized contact counts. Each dot in the plot corresponds to a particular genomic distance. In each case, we observe strong overdispersion (i.e., points are above the diagonal), supporting the idea to model Hi-C count data with overdispersed models.

**Fig. 1.**
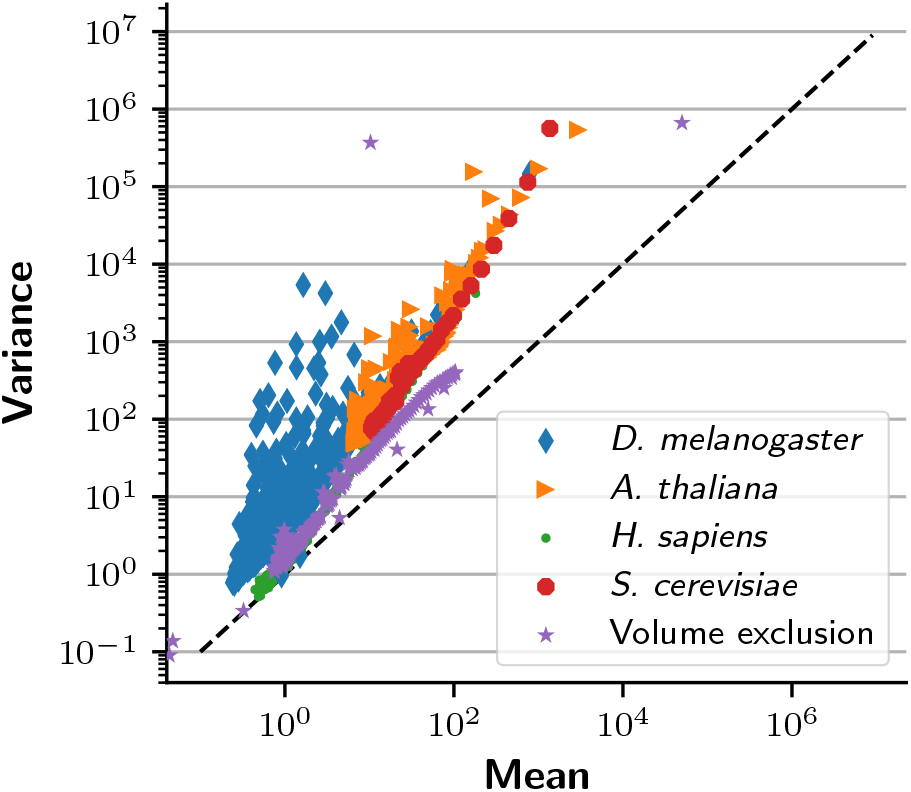
Mean and variance of contact counts in different Hi-C datasets. Each point represents a given genomic distance in one dataset, where we estimate the mean and variance of contact counts. The dashed line corresponds to the relationship assumed by the Poisson model: *σ*^2^ = *µ*. The “Volume exclusion” dataset is simulated following a previously described model^35^.

We then repeat this experiment on simulated data by generating 50,000 structures using a previously described volume exclusion model of the budding yeast^35^. From this population of structures we create a contact count matrix at 3 kb resolution, assuming that loci closer than 40 nm come into contact. This simulated contact count map has been shown to be highly correlated with experimental Hi-C data^35^. The resulting dataset (purple series in Figure 1) displays the same overdispersion as the data. We thus conclude that the overdispersion is an inherent property of Hi-C data and not an experimental artifact. We hypothesize that the overdispersion arises due to the large variety of different structures present in a single Hi-C experiment.

### IV.B. Pastis-NB is accurate and robust on simulated data

Next, we use simulated data to compare our approach, which we refer to as Pastis-NB, with four different algorithms. Pastis-MDS is a weighted metric MDS method that attempts to place the beads such that the distance between each pair matches as closely as possible the wish distances derived from contact counts^36^. ChromSDE is a variant of metric MDS that penalizes non-interacting pairs of beads to keep them far away from one another and optimizes the counts-to-distances mapping coefficient with a golden search^39^. ShRec3D is a two-step method that first derives distances from contact counts using a shortest path algorithm and then applies classical MDS^19^. Pastis-PM models contact counts as Poisson random variables with the 3D structure as a latent variable and casts the inference problem as a likelihood maximization, optimizing the jointly the structure and the parameters of the counts-to-distance function. The five methods fall into two categories: chromSDE, Pastis-PM, and Pastis-NB are *non-metric* because they fit a parametric curve to estimate a count-to-distance mapping from the data; while Pastis-MDS, and ShRec2D are *metric* because they do not do this fitting. In our experiments, each software package is used with default parameters.

#### IV.B.1. Robustness to coverage and dispersion

We first run all algorithms on the datasets with varying dispersion and coverage. Our goal here is to assess how well the different methods reconstruct a known 3D structure from simulated data at different coverage and dispersion levels. High coverage typically corresponds to a high signal-to-noise ratio, whereas low coverage yields sparse, low signal-to-noise ratio matrices. Similarly, when the dispersion parameter tends to infinity, the negative binomial distribution (by definition overdispersed) tends to a Poisson with lower variance. Thus, the lower the dispersion parameter, the noiser the dataset. All methods should therefore perform better as the dispersion parameter (*γ* in our setting) and the coverage increase.

In this first series of experiments, we provide the correct count-to-distance or distance-to-count transfer functions to the metric methods, who need it. In this setting, for infinite coverage and infinite dispersion parameter, all methods should consistently estimate the correct structure, at least if they manage to converge to the global optimum of their objective function.

We first plot the average root mean squared deviation (RMSD) error between the true and predicted structures, as a function of coverage (Figure 2A) for the datasets with varying coverage. Strikingly, ShRec3D’s and Pastis-NB’s results are extremely stable to coverage, while the three other methods see their performance decrease with coverage. ShRec3D performs relatively well when the coverage is low but poorly in the high coverage setting. All the other methods perform similarly well for high coverage, with Pastis-PM and Poisson-NB achieving the lowest RMSD at high coverage, but exhibit strong differences in the low coverage setting. In the low coverage setting, Pastis-NB remains extremely good, with barely any decrease in performance even at the lowest coverage, while Pastis-MDS’ performance degrades quickly with less than 70% of the coverage, and similarly Pastis-PM and chromSDE see their performance deteriorate with less than 40% of the coverage. With the best RMSD error among all methods for all coverage, Pastis-NB is the clear winner in this experiment.

**Fig. 2.**
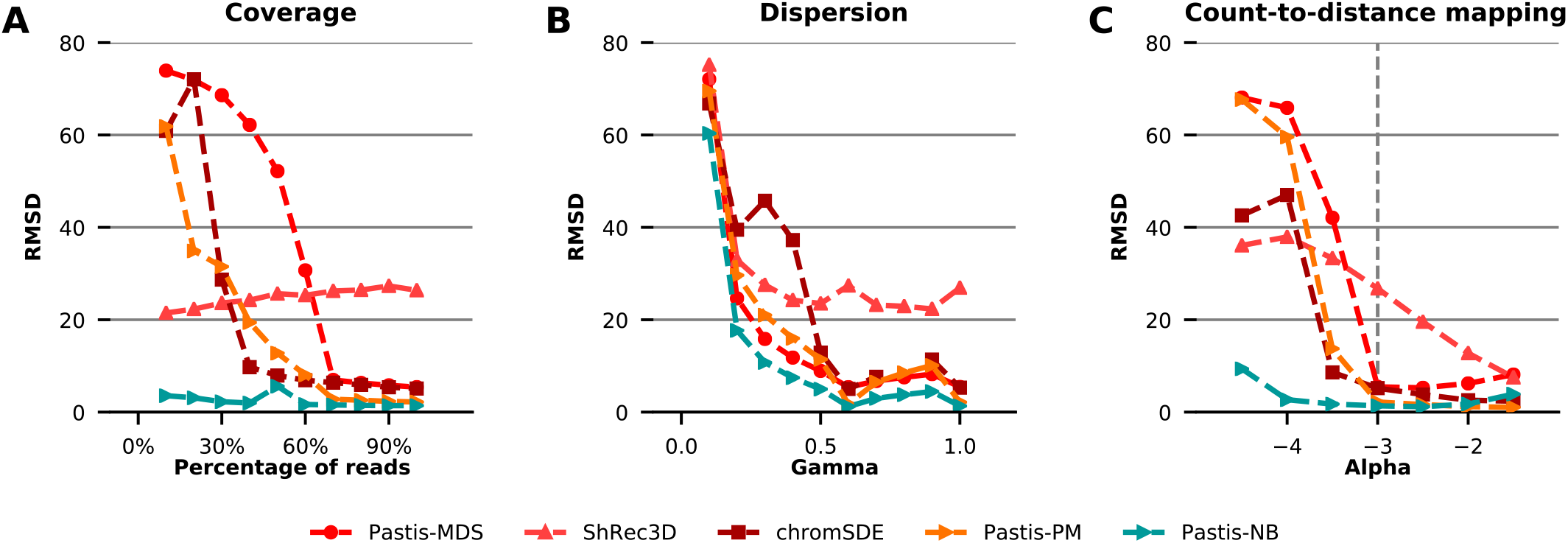
Performance evaluation on simulated data. Each plot shows the mean RMSD error (over 10 random simulations with different random seed) between the predicted structure and the true structure, for five different methods, when one parameter of the simulationis varied. **A**. The parameter *β* is varied such that the coverage is 10–100% of the original dataset. **B**. The parameter *γ*, which controls over-dispersion, is varied between 0.1 and 1. Smaller values correspond to more over-dispersion. **C**. The paramater *α*, which controls the count-to-distance mapping coefficient, is varied between −4.5 and −1.5.

We then plot the average RMSD as a function of the dispersion tuning parameter *γ* (Figure 2B) for the second set of simulated contact maps with varying dispersion. As expected, we observe that all methods tend to perform better when *γ* increases (corresponding to less overdispersion), and have poorer performance when the data are too overdispersed. ShRec3D’s results tend to be stable to changes in dispersion, but worse than other methods for large *γ*’s. All methods perform poorly in the highest dispersion setting. ChromSDE performs relatively poorly in the medium to high dispersion setting. Pastis-NB has again the best performance across all dispersion values, although the difference relative to other methods, particularly Pastis-PM and Pastis-MDS, is small.

#### IV.B.2. Robustness to incorrect parameter estimation

We then compare the algorithms on datasets with varying counts-to-distances mappings. Metric methods (Pastis-MDS and ShRec3D) require as input a count-to-distance transfer function. While Pastis-MDS relies on ideal physical laws to define this mapping, ShRec3D uses *ad hoc* conversion of counts into physical distances. However, DNA may not follow the ideal properties of polymers underlying the default transfer function; thus, structures inferred from these methods may diverge from the correct ones. Our goal here is to assess how well the different methods perform when the transfer function is mis-specified. We expect non-metric methods to perform better on these datasets, because they should be able to adapt the transfer function to best fit the data.

Figure 2C shows the average RMSD error of each algorithm as a function of the *α* parameter used to simulate the data. It is worth noting that the lower the *α* parameter is, the noisier the simulated contact map is: a low *α* parameter indeed results in a contact count map with very few long range interaction counts.

Pastis-NB works well across all values of *α*, exhibiting a striking difference from the rest of the methods for *α*≤ − 3. Notably, all methods perform much better for high *α* than for low *α*. This phenomenon can be explained by the properties of the contact count maps in this setting: low *α* values in the count-to-distance function lead to abrupt changes in the probability of seeing contact counts between small and large distances, whereas high *α* values yield a much more uniform expected contact counts map. Thus, for identical coverage, low *α* datasets are much sparser than high *α* datasets. In short, despite identical coverage in all datasets, the signal-to-noise ratio varies strongly with *α*, thereby leading to much better overall performance for low *α* both for metric and non-metric methods, even when the transfer function is mis-specified.

#### IV.C. Pastis-NB yields stable and consistent structures on real Hi-C data

We now test the different methods on real Hi-C data. Since in this case the true consensus structure is unknown, we investigate instead the behaviors of the different methods in terms of their ability to infer consistent structures from replicate datasets and across resolutions.

#### IV.C.1. Pastis-NB shows increased stability when performing the inference without filtering zero counts

Before delving into a detailed comparison of Pastis-NB with other methods on various tasks, we first illustrate in this section the benefits brought by including zero-valued counts in the model inferred by Pastis-NB. This is to be contrasted with many previously published methods that perform 3D structure inference using only a subset of the data, and in particular disregard zero counts. For example, Tanizawa *et al*. ^34^ and Duan *et al*. ^10^ consider only the top 2% significant contact counts, whereas Varoquaux *et al*. ^36^ exclude zero contact counts from the inference. Many MDS-based methods require a transformation of contact counts into distances: this is often based on a power-law relationship with a negative coefficient and is thus undefined for zero contact counts. As a result, methods include zero contact counts either through an *ad hoc* penalization term on non-interacting beads or by converting zero counts to an *ad hoc* distance (for example, the largest distance obtained on non-zero counts or using prior knowledge of the structure). In contrast, Pastis-NB formulates the inference in a way that naturally includes zero contact counts.

To assess the impact of zero counts information, we compare the default Pastis-NB model, which takes into account zero counts in the likelihood objective function (2), with a variant where we only retain non-zero counts. We assess the stability of the inferred structures across biological replicates when using all counts versus only non-zero counts. To do so, we run the inference on the whole sets of contact counts versus the filtered one, on both replicates 75 and 76 of KBM7, for contact count matrices at five different resolutions. We thus infer two structures for each autosomal chromosome, one for each replicate. We then rescale them such that the structures fit in a sphere of a predefined diameter, and we compute the RMSD between the two rescaled structures as described in Varoquaux *et al*. ^36^, to estimate the stability of the inference. Recall that the different methods cannot optimize the coordinates of beads that have zero contact counts. Thus, before computing the RMSD, we filter out from both structures any beads that have zero contact counts in either dataset.

We then compare the stability of the inference between the two approaches (filtered versus unfiltered zero counts). Figure 3A shows, for each approach, the distribution of RMSD values across the 22 chromosomes and five resolutions. We clearly see that keeping all zero counts leads to significantly smaller RMSD, hence more stable structures across biological replicates.

**Fig. 3.**
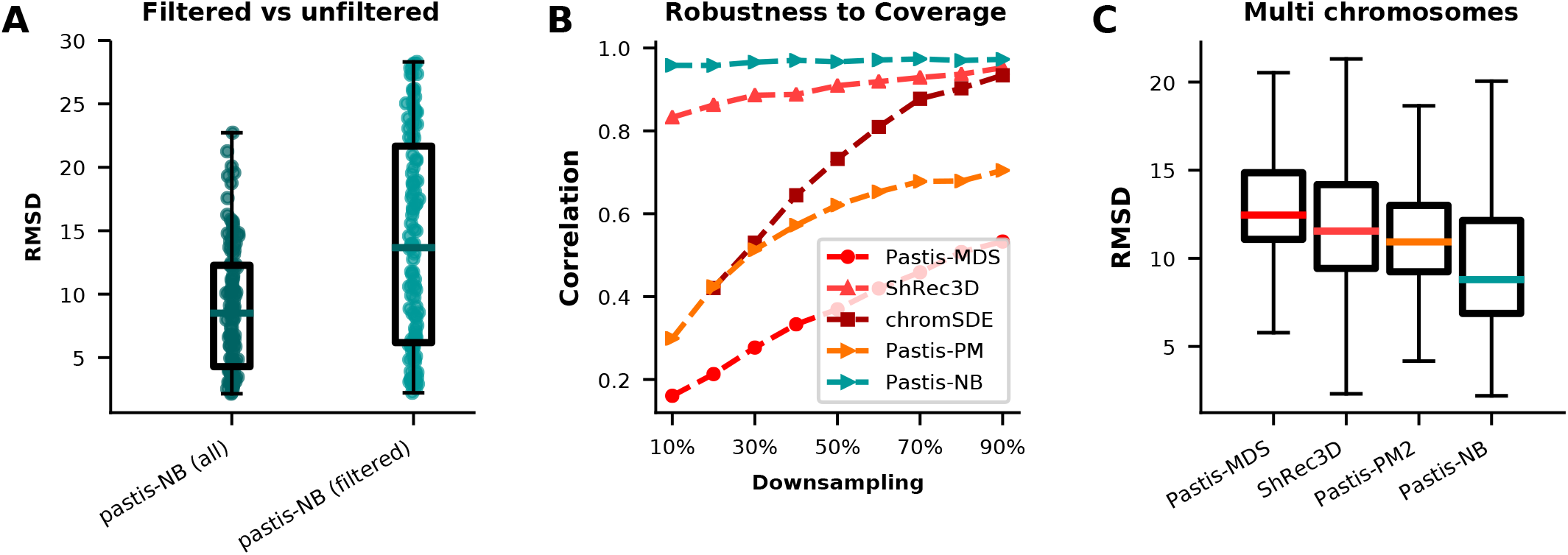
Stability results on Rao *et al*. ^*25*^ and Duan *et al*. ^*10*^A. The RMSD between pairs of structures inferred by Pastis-NB using all contact counts (“all”) or excluding zero contact counts (“filtered”). **B**. Inference is performed on a downsampled contact map and the resulting structure is compared to the structure obtained using the whole dataset. ChromSDE fails to infer a structure with the correct number of bins on the datasets downsampled to 10% of the original coverage: results for chromSDE at 10% are thus not displayed. **C**. The RMSD between pairs of chromosomes inferred by subsampling chromosomes from^10^‘s *S. cerevisiae* dataset.

#### IV.C.2. Stability to replicates

Replicates, which involve multiple runs of the same experiment performed on similar samples with the same experimental settings, are typically carried out to assess variability of the results. Because the underlying 3D model should not change in this case, we compare the results of the inference of the different algorithms on two replicates of the nearly haploid human KBM7 cell line. Similar to what we did in Section IV.C.1, we infer two structures on the two replicates of interest for each autosomal chromosomes. We compute the RMSD between each pairs of structures as described above, as well as the Spearman correlation of the distances.

Not surprisingly, the stability of the inference across replicates decreases as the resolution grows in all methods. Table II shows the average correlation reached by each method at each resolution, and Supplementary Table S2 shows the corresponding average RMSD. At low resolution (1 Mb, 500 kb), chromSDE performs the best both in terms of correlation and in terms of RMSD between replicates, although all methods perform well in that setting with correlations ranging between 0.95 and 0.99. At high resolution (250 kb, 100 kb, 50 kb), Pastis-NB performs the best, both in correlation and in RMSD, despite the non-convex nature of the optimization problem solved. It is remarkable that even at 50 kb, Pastis-NB reaches a correlation of 0.96 between replicates, while the second best method (ShRec3D) see its correlation decrease to 0.87, and chromSDE, which is the most stable at low resolution, only obtains a correlation of 0.62.

**Table II.** Stability across replicates. The table shows the average Spearman correlation between structures inferred on biological replicates on 22 autosomes of the KBM7 cell line at varying resolutions. In bold is the best correlation per row.

|  | Pastis-MDS | ShRec3D | chromSDE | Pastis-PM | Pastis-NB |
| --- | --- | --- | --- | --- | --- |
| 1mb | 0.95 | 0.97 | **0.99** | 0.96 | 0.96 |
| 500kb | 0.97 | 0.96 | **0.98** | 0.95 | 0.97 |
| 250kb | 0.92 | 0.93 | 0.97 | 0.89 | **0.98** |
| 100kb | 0.59 | 0.89 | 0.87 | 0.73 | **0.97** |
| 50kb | 0.33 | 0.87 | 0.62 | 0.56 | **0.96** |

#### IV.C.3. Stability to resolution

Zhang *et al*. ^38^ show that the mapping from contact counts to physical distance differs from one resolution to another, underscoring the importance of good parameter estimation. To study the stability of the structure inference methods to changes in resolution, we compute the RMSD between pairs of structures inferred at different resolutions (1 Mb, 500 kb, 250 kb, 100 kb, and 50 kb). Each inferred structure is rescaled such that all beads fit in a nucleus of size 100. To compare two structures at different resolutions, we downsample the structure of the highest resolution by averaging its coordinates until it is of the same resolution as the other one. We then compute the RMSD and correlation between the two structures of the same size.

Results of this experiment (Table III) show that Pastis NB is more stable to resolution changes than other methods, both in terms of RMSD and in terms of correlation, with for example a decrease of ∼20% in average RMSD compared to ShRec3D, the second best (14.50 vs 18.27).

**Table III.** Stability across resolution. The table lists the average RMSD and Spearman correlation between pairs of structures for replicate 75 at different resolutions. In bold are the lowest average RMSD and highest average Spearman correlation. These values were computed on the KBM7 nearly haploid human cell line^25^.

|  | Pastis-MDS | ShRec3D | chromSDE | Pastis-PM | Pastis-NB |
| --- | --- | --- | --- | --- | --- |
| RMSD | 26.57 | 18.27 | 21.08 | 22.25 | **14.50** |
| Correlation | 0.48 | 0.79 | 0.65 | 0.66 | **0.86** |

#### IV.C.4. Stability to coverage

We then study the stability of the structure inference methods to coverage. To do so, we downsample the 100 kb contact count matrices between 10% and 90% of the original coverage. We perform inference from these downsampled contact maps and compute the Spearman correlation between Euclidean distances of the obtained structures and the structures inferred on the full matrix. Results of this experiment (Figure 3B) show that all methods tend to see the correlation decrease with the downsampling, as expected. While ShRec3D and chromSDE yield high correlations at high coverage, the correlation decreases sharply for chromSDE, reaching ∼0.3 at 10% downsampling, and a bit less sharply for ShRec3D, which reaches ∼0.8 at 10% downsampling. Pastis-PM and Pastis-MDS have the worst correlation at high coverage, and see their correlation decrease sharply with coverage. Pastis-NB stands out as the method with largest correlation at high coverage, but also as the method that witnesses barely any decrease in correlation when coverage decreases.

#### IV.C.5. Whole genome inference

Finally, we consider the harder task of whole genome inference, rather than inferring structures separately per chromosome. When tackling whole genome inference, a new problem arises for non-metric methods: the *inter* - chromosomal contact counts dominate the estimation of the counts-to-distance parameter *α*. Indeed, while the estimation of the *α* parameter is very stable on single chromosomes, we observe that this parameter *α* increases to −1 when the number of chromosomes increases for some non-metric methods. This has the effect of collapsing beads belonging to a chromosome together, while pushing beads belonging to other chromosomes away from one another.

To assess how well whole genome inference performs, we perform yet another stability experiment. We randomly subsample the number of chromosomes of a *S. cerevisiae* dataset and perform the 3D structure inference. We then assess the stability of the structure inference by computing the RMSD and Spearman correlation for each chromosome independently and taking the average to obtain a single RMSD and Spearman correlation score. Note that chromSDE does not support multi-chromosome inference: we have thus excluded chromSDE from these results. Results once again show that Pastis-NB is the most stable method both in RMSD (Fig 3C) and in correlation (Supp. Fig. S1).

## V. DISCUSSION AND CONCLUSION

We present in this work a new model, Pastis-NB, to infer a consensus 3D structure of the genome from Hi-C contact count data. We model interaction counts as negative binomial random variables, and we cast the inference as a likelihood maximization problem. Modeling counts as negative binomial random variables allows us to better model the presence of overdispersion in Hi-C data, which we observed experimentally in Hi-C data from different organisms. Through extensive experiments on simulated and real Hi-C data, we showed that Pastis-NB consistently outperforms a representative set of four competitive methods across a range of metrics. In particular, Pastis-NB yields remarkably stable and accurate results in the case of highly dispersed contact count data. This improvement is particularly striking at high resolution and at low coverage, with 3D models inferred much more robustly with Pastis-NB than with other methods.

A limitation of Pastis-NB resides in the inference consensus 3D models of chromosome architecture. Consensus models are not necessarily representative of the true folding of DNA in the cell. For example, a consensus model of *S. cerevisiae*’s genome will sometimes cluster telomeres together, while it is known that telomeres tether at the nuclear membrane. One should thus interpret these models with care. Yet consensus methods are powerful dimensionality reduction tools that can serve as an entry point for many analyses, such as data integration and visualization.

## VI. ACKNOWLEDGEMENTS

This work was supported by NIH awards U54 DK107979 and UM1 HG011531.

