## Supplementary data for "Inference of genome 3D architecture by modeling overdispersion of Hi-C data"

### Supplementaries materials

January 29, 2021

#### List of Tables


1 **Percentage of loci filtered by dataset prior to normalization** . . . . . . . . . . . . . . . 3

2 **Stability across replicates** The table shows the average RMSD between structures inferred on biological replicates on 22 autosomes of the KBM7 cell line at 1 Mb, 500 kb, 250 kb, 100 kb, and 50 kb. In bold is the best average RMSD value. . . . . . . . . . . . . . . . . . . . . . . . . 4


#### List of Figures


1 **Multi-chromosomes structure inference.** The Spearman correlation between pairs of chromosomes inferred by subsampling chromosomes from Duan et al. [2010]'s *S. cerevisiae* dataset. . . . . . . . . . . . . . . . . . . . . . . . . . . . . . . . . . . . . . . . . . . . . . . 2


#### Contents


**1 Supplementary Figures** **2**

**2 Supplementary tables** **3**

**3 Supplementary materials** **4**

3.1 Normalization of the data . . . . . . . . . . . . . . . . . . . . . . . . . . . . . . . . . . . . . 4

### 1 Supplementary Figures

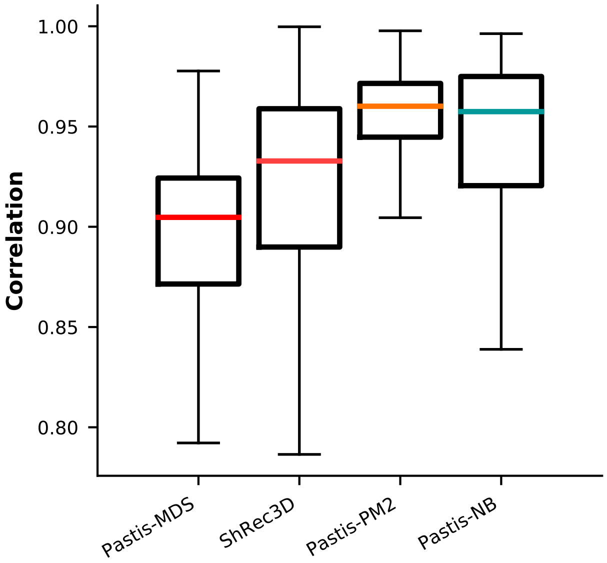


**Supplementary Figure 1: Multi-chromosomes structure inference. The Spearman correlation between pairs of chromosomes inferred by subsampling chromosomes from Duan et al. [2010]'s *S. cerevisiae* dataset.**

### 2 Supplementary tables

| **Dataset** | **% loci filtered** |
|---|---|
| Rao et al. [2014] (1 mb) | 1% |
| Rao et al. [2014] (500 kb) | 2% |
| Rao et al. [2014] (250 kb) | 3% |
| Rao et al. [2014] (200 kb) | 4% |
| Rao et al. [2014] (100 kb) | 5% |
| Rao et al. [2014] (50 kb) | 6% |
| Duan et al. [2010] (10 kb) | 4% |
| Sexton et al. [2012] (10 kb) | 4% |
| Feng et al. [2014] (40 kb) | 4% |

**Supplementary Table 1: Percentage of loci filtered by dataset prior to normalization**

| | pastis-MDS | ShRec3D | chromSDE | pastis-PM2 | pastis-NB |
|---|---|---|---|---|---|
| 1mb | 9.99 | 6.53 | **5.09** | 10.45 | 9.61 |
| 500kb | 8.64 | 9.65 | **7.17** | 11.09 | 8.25 |
| 250kb | 11.65 | 12.71 | 8.07 | 16.04 | **7.96** |
| 100kb | 24.38 | 14.63 | 13.91 | 21.49 | **10.02** |
| 50kb | 31.29 | 14.62 | 23.35 | 26.16 | **10.60** |

**Supplementary Table 2: Stability across replicates The table shows the average RMSD between structures inferred on biological replicates on 22 autosomes of the KBM7 cell line at 1 Mb, 500 kb, 250 kb, 100 kb, and 50 kb. In bold is the best average RMSD value.**

### 3 Supplementary materials

#### 3.1 Normalization of the data

The raw contact count matrix **c** suffers from many biases, some technical (from the sequencing and mapping) and others biological (inherent in the physical properties of chromatin) [Imakaev et al., 2012, Yaffe and Tanay, 2011]. Imakaev et al. [2012] proposed a simple iterative correction and eigenvalue decomposition method called ICE to estimate the biases and normalize the data. This method relies on two primary assumptions. First, the biases of each entry $c_{ij}$ can be written as the product of two biases $b_i$ and $b_j$ associated with loci $i$ and $j$; thus, if $\mathbf{c}^N$ is the normalized contact count matrix, then $c_{ij} = b_i b_j c_{ij}^N$. Second, the total number of contacts associated with each locus should be equal. From these assumptions, we can formulate a non-convex optimization problem to estimate the vector of biases $b$. The problem can be solved exactly using an iterative procedure akin to the Sinkhorn algorithm. We apply the ICE method to all of the data used in this study. Prior to normalization, we filter out rows and columns having the fewest number of contacts to avoid degeneracies during the iterative correction (see Supplementary Table 1).
